## Supplementary figures + variant models for "Self-inhibiting percolation and viral spreading in epithelial tissue"

### I. EMPIRICAL DATA

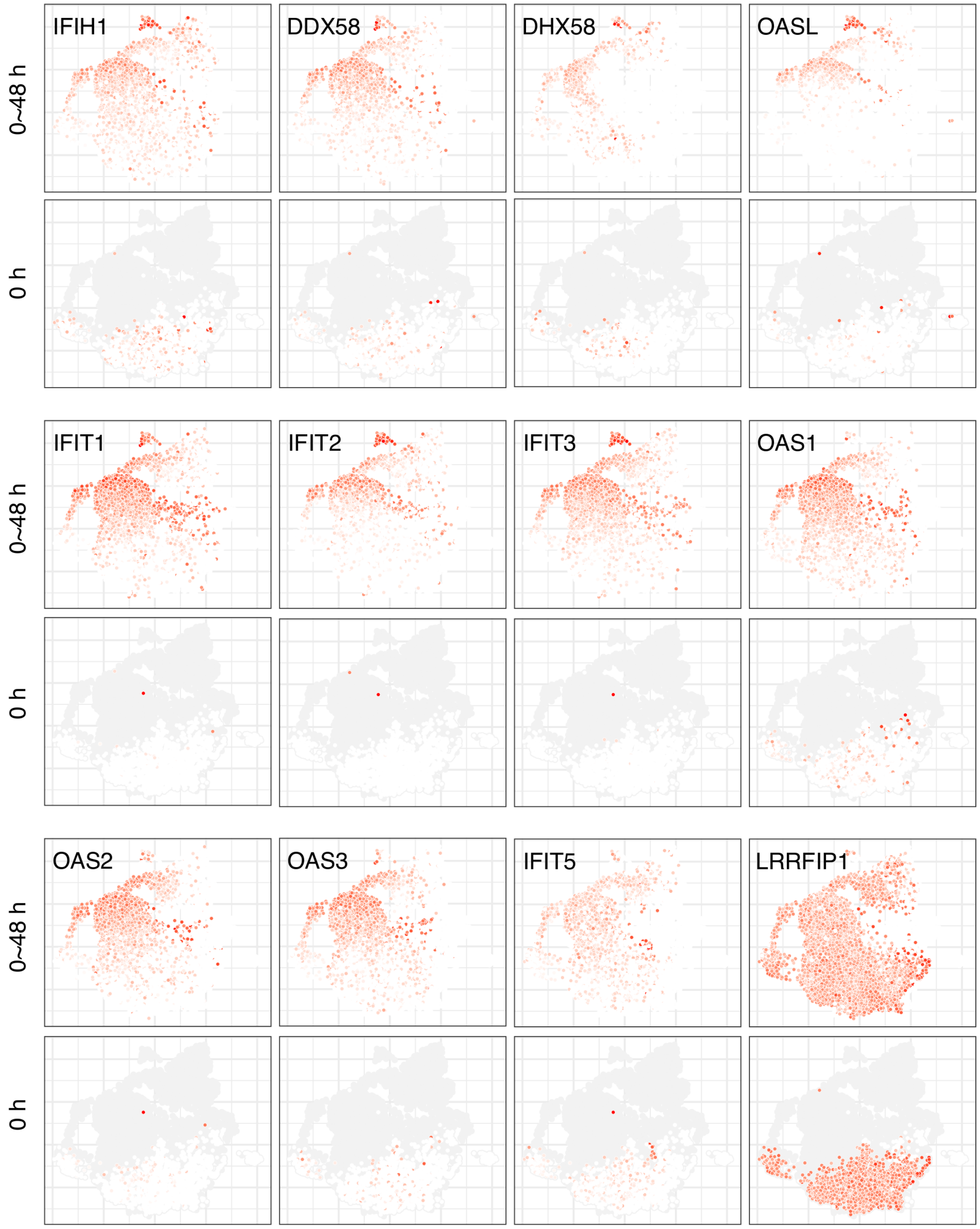


FIG. S1. Expression of antiviral genes at different stages of early SARS-CoV-2 infection. Once the infection has proceeded for two days, cells in the $A$ state express all the antiviral genes highly. The typical $a$ state can be identified from the 0 h panels where some cells express low to moderate levels of antiviral genes before any exposure to infection. 8% of the cells at 0 h express at least two of IFIH1, DDX58 and DHX58 at a detectable level.

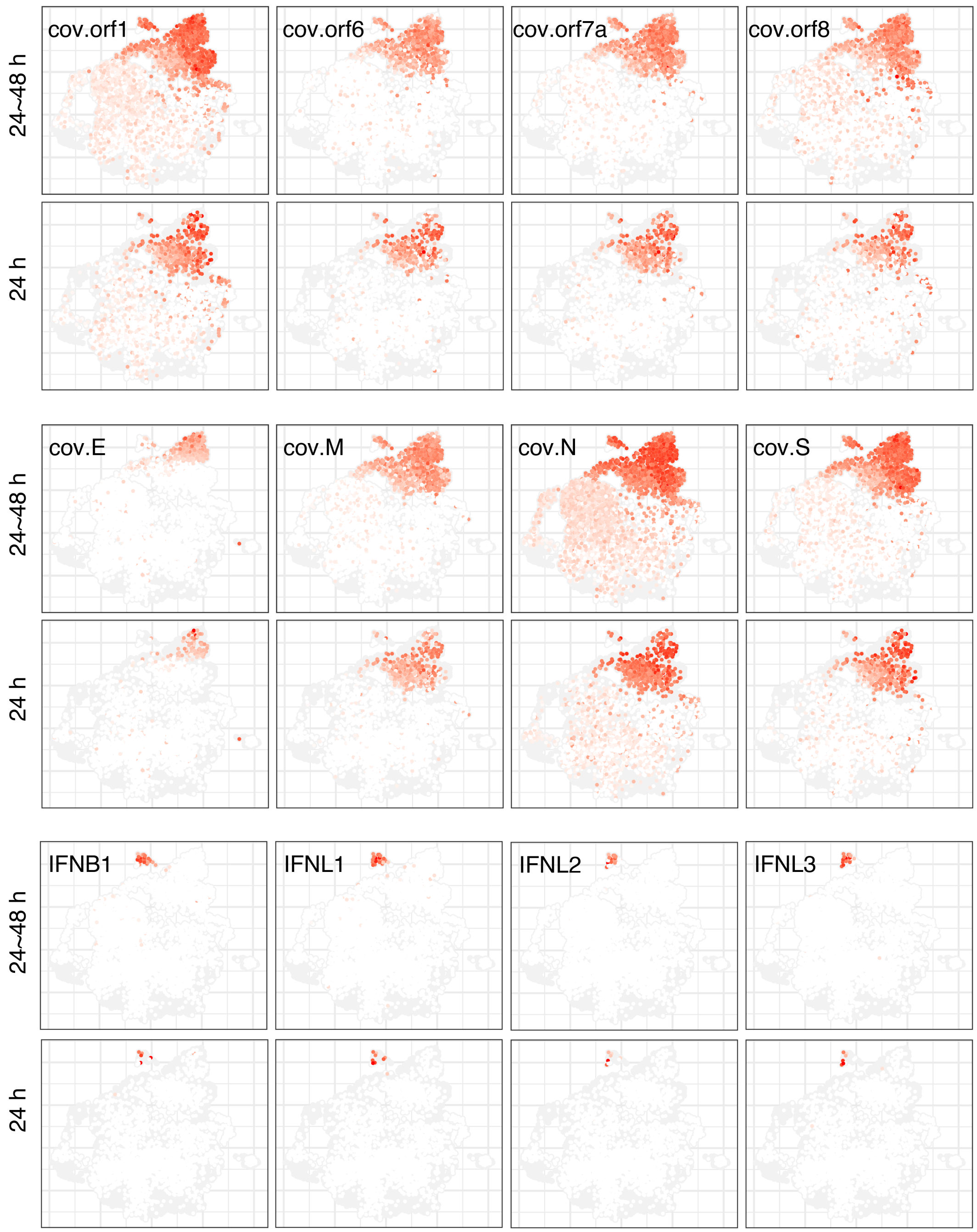


FIG. S2. Detected viral genes and expression of IFNs at different stages of early SARS-CoV-2 infection. Once the infection has proceeded for two days, viral genes (cov.*) are detected at a high level in the $V$ cells. The IFN genes are mainly expressed in $N$ cells, amounting to 0.6% at 24h and to 4.8% at 48h.

### II. THE OVA MODEL

Fig S3 introduces the simplified OVA model, which does not include an $N$ state, but rather gives each susceptible cell a probability of spontaneously converting to an antiviral state in each time step.

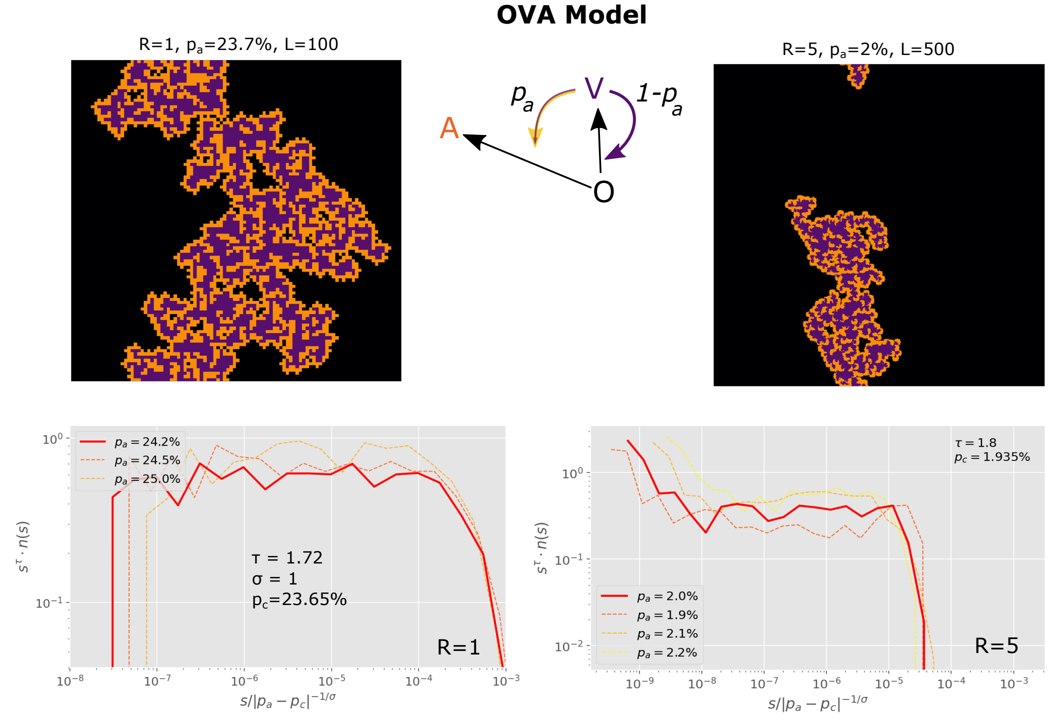


FIG. S3. OVA Model: In each time steps, $L^2$ updates are made. At each update, one of the $L \times L$ sites is randomly selected. If this site is in the $O$ state, nothing happens. If the site is $V$ state then its nearest 4 neighbors are exposed to the virus, given that they are susceptible (i.e. in the $O$ state). One then in random order selects each of the neighbors that are in $O-$ state. For each selected neighbor $i$ one draws a random number $ran_i \in [0, 1]$. If $ran_i \geq p_A$ the neighbor is flipped to $V$ state. If, on the other hand, $ran < p_a$ then one converts all $O$ cells within radius $R$ around the neighbor $i$ into the $A$ state. These changes include the neighbor $i$ itself.

### III. TIME-EVOLUTION OF STATE OCCUPANCY

In Fig. S4, we show the time-evolution of occupation fractions for the different states of the model, for various values of $p_a$ below the critical value $p_c$, for two interferon spreading radii, $R = 1$ and $R = 5$. Each panel is based on a single typical realization.

As shown qualitatively in the figure, the speed of propagation as well as the final occupancy ratios depend on the distance to the threshold, $|p_a - p_c|$.

### IV. STOCHASTIC CONVERSION

While even the base model has a level of stochasticity – since $L^2$ are randomly chosen, with replacement, to be updated in each time step – we here simulate a version of the dynamics which includes stochastic conversion, i.e. each action of a cell on a neighboring cell occurs only with a probability $p_{conv}$ (and the original model is recovered as the $p_{conv} = 1$ scenario). This necessarily slows down the dynamics (or effectively rescales time by a factor $p_{conv}$), but crucially we find that it does not appreciably affect the location of the threshold $p_c$. In Fig. S5, we show a parameter scan across $p_a$ values for $R = 1$ and $p_{conv} = 0.5$, which shows that the threshold continues to exist at around $p_a = 27\%$.

### V. SIMULATIONS WITH A GAUSSIAN KERNEL

In the NOVAa model of the main text, the spread of interferons (i.e. the action of cells in the $N$ state) always follows a circular motif. When an $N$ cell is selected, it will act on all cells within a radius $R$ (provided they are in the $O$ or $a$ state). To more closely approximate the diffusion of interferons - and to allow for some stochasticity in this

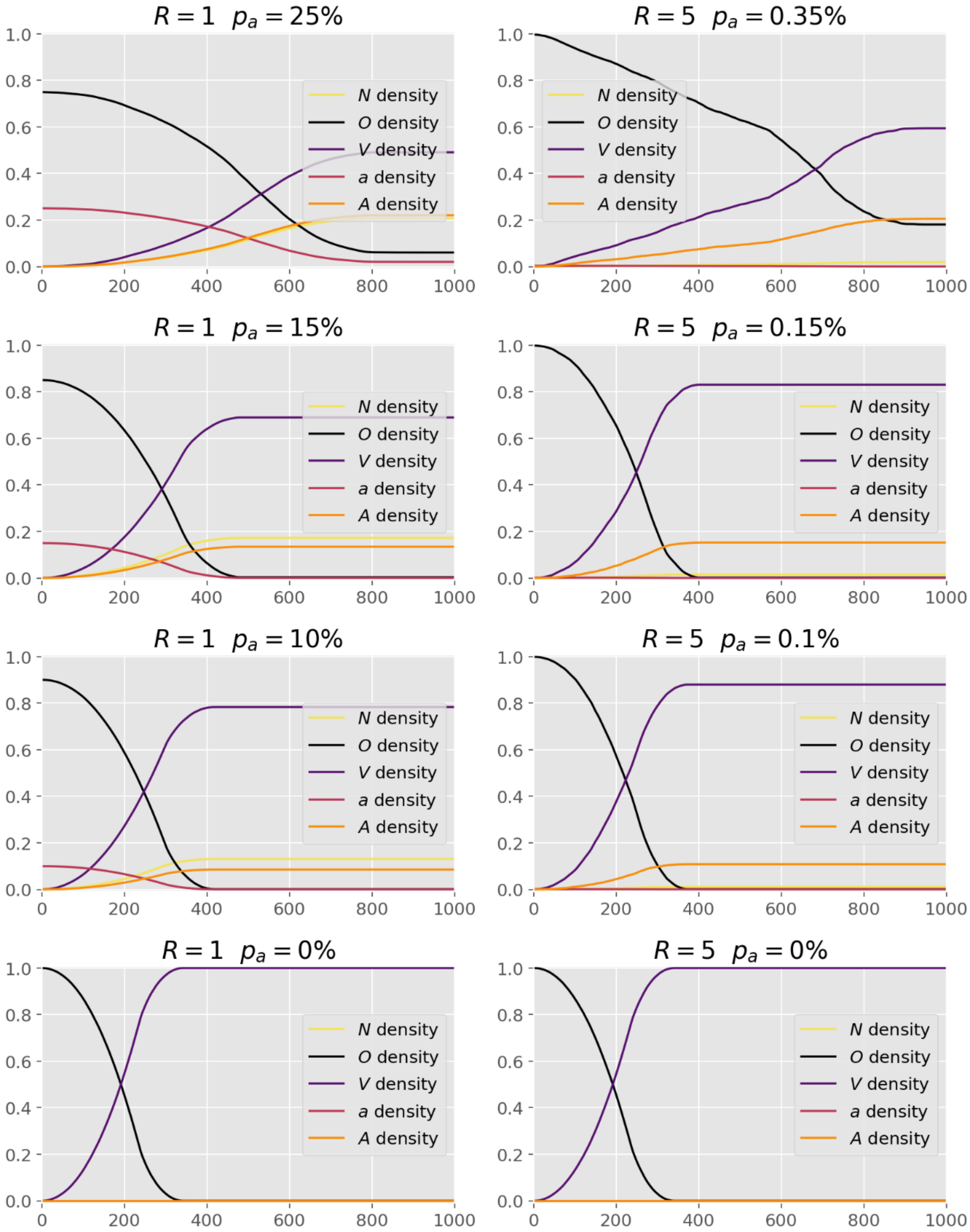


FIG. S4. Cell fractions within the different states over times, at a range of $p_a$ values (below the critical value $p_c$), for two interferon spreading radii, $R = 1$ and $R = 5$.

process - we will here consider an extension of the model, in which the spread of interferon is modeled by a Gaussian kernel.
In the following, we will refer to the model presented in the main text as the *model with a circular spreading motif* and the alternative model as the *Gaussian model*.
The Gaussian model is implemented as follows:
Let

$$P(d;\sigma) = \mathcal{N}\exp\left[-d^2/(2\sigma^2)\right] \tag{1}$$

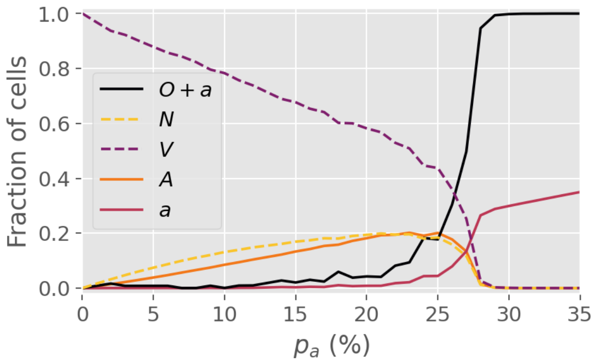


FIG. S5. **Stochastic conversion.** Average final-state cell fractions in a variant of the model with stochastic conversion, i.e. in which each conversion of a neighboring cell takes place with a probability $p_{conv} < 1$. In this figure, $p_{conv} = 0.5$. Note that the critical threshold is largely unaffected. The figure is based on 4000 simulations.

with $\mathcal{N}$ a normalization constant and $\sigma = \frac{2\sqrt{2}}{3\sqrt{\pi}}R$. This value of $\sigma$ ensures that the mean distance to converted cells (i.e. those acted upon by $N$ cells) is the same as in the model with a circular spreading motif.
The normalizaton constant $\mathcal{N}$ is chosen such that the average number of converted cells is the same as in the model with a circular spreading motif. This results in a value of $\mathcal{N} = \mathcal{N}_R/(2\pi\sigma^2)$ where $\mathcal{N}_R$ is the number of lattice points within a radius of $R$ from a central lattice point.

The simulation routine then proceeds as follows:

- At each time step, $L^2$ random sites are selected for updating (with replacement).
- For non-$N$ cells, updates are carried out as in the main text.
- When an $N$ cell is selected, the update proceeds as follows:
  - All cells within a radius of $4\sigma$ are designated as neighbors.
  - For each of these neighbor cells, convert the cell (i.e. let the $N$ cell act on it) with probability $P(d;\sigma)$, where $d$ is the Euclidean distance between the neighbor cell and the $N$ cell.
  - Once all neighbours have been considered, move the $N$ cell to a new state $N_1$
- When an $N_1$ cell is selected, it behaves identically to an $N$ cell. Once an $N_1$ cell has been updated, it moves to state $N_2$, which is inactive.

The extended radius of $4\sigma$ was chosen to ensure that the vast majority of potential interactions are included, while retaining numerical efficiency.

The introduction of the two new (albeit very simple) states $N_1$ and $N_2$ was to ensure that a single $N$ cell does not act on a very large number of other cells simply by being selected multiple times during a simulation. In the circular motif case, this was automatic since an $N$ cell could only act on the same $\mathcal{N}_R$ cells in each time step, and once they were in the $N$, $V$ or $A$ state, the $N$ cell could no longer affect them. In practice, an $N$ cell could act twice on a susceptible cell, once to turn it from $O$ to $a$ and once to convert it from $a$ to $A$.

In the Gaussian case, an $N$ cell could in principle act on an unlimited number of cells, although the rate would decrease with distance according to the Gaussian kernel. Thus, it was necessary to introduce some memory in the form of the $N_1$ and $N_2$ states to more closely mimic the circular motif case of the main text.

As shown in Fig S6, the Gaussian model can behave quite differently to the circular motif model even given the same parameter values. The primary reason for this difference owes to the nonzero probability of long-range conversions in the Gaussian model, since this allows for bridging areas otherwise devoid of $a$ cells.

### VI. CODE AVAILABILITY

The C++ source code for the simulations and a Python notebook for plotting can be found at:
`https://github.com/BjarkeFN/ViralPercolation`

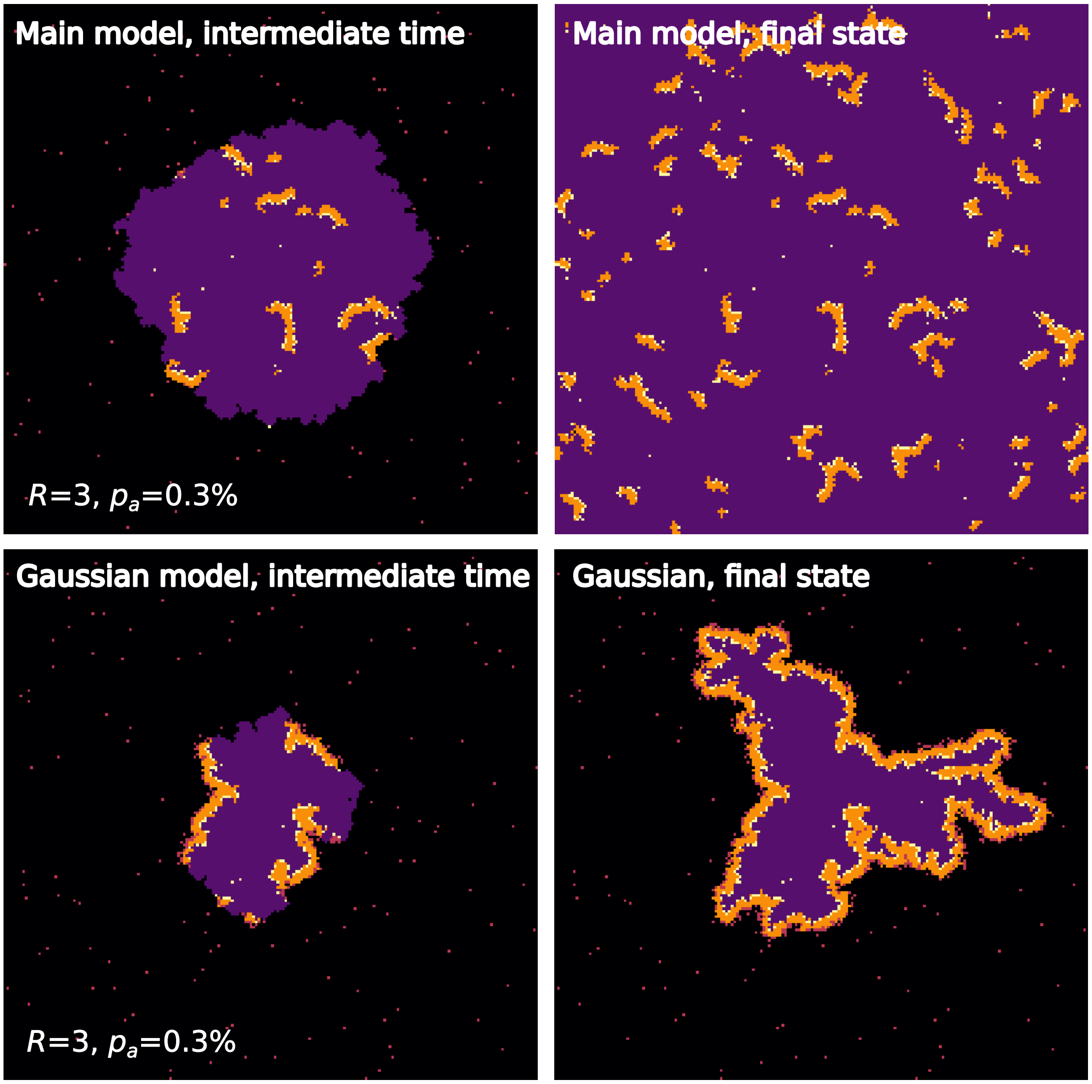


FIG. S6. Comparison of the Gaussian Model and the main NOVAa model at identical parameters.
